# PCBP1 Deficient Pigs Hold the Potential to Inhibit CSFV Infection

**DOI:** 10.1101/2021.12.23.474075

**Authors:** Chunyun Qi, Daxin Pang, Kang Yang, Shuyu Jiao, Heyong Wu, Chuheng Zhao, Lanxin Hu, Feng Li, Jian Zhou, Lin Yang, Dongmei Lv, Xiaochun Tang, Hongsheng Ouyang, Zicong Xie

**Affiliations:** Key Lab for Zoonoses Research, Ministry of Education, Animal Genome Editing Technology Innovation Center, Jilin Province, College of Animal Sciences, Jilin University, Changchun130062, China; Chongqing Research Institute, Jilin University, Chongqing401123, China; Chongqing Jitang Biotechnology Research Institute Co., Ltd; Shenzhen Kingsino Technology Co., Ltd., Shenzhen, China

## Abstract

Classical swine fever virus (CSFV), pathogen of classic swine fever, has caused severe economic losses worldwide. Poly (rC)-binding protein 1 (PCBP1), interacting with N^pro^ of CSFV, plays a vital role in CSFV growth. Here, our research is the first report to generate PCBP1 knockout pigs via gene editing technology. The PCBP1 knockout pigs exhibited normal birth weight, reproductive-performance traits, and developed normally. Viral challenge results indicated that primary cells isolated from F_0_ and F_1_ generation pigs could significantly reduce CSFV infection. Additional mechanism exploration further confirmed that PCBP1 KO mediated antiviral effect is related with the activation of type I interferon. Beyond showing that gene editing strategy can be used to generate PCBP1 KO pigs, our study introduces a valuable animal model for further investigating infection mechanisms of CSFV that help to develop better antiviral solution.

**Importance:** As a negative regulator in immune modulation, the effects of PCBP1 on viral replication have been found to be valuable. Here, this study was the first report to generate PCBP1 knockout pigs with normal pregnancy rate and viability. Primary cells isolated from F_0_ and F_1_ generation PCBP1 knockout pigs could significantly reduce CSFV infection. The PCBP1 knockout pigs could be used as a natural host models for investigating the effects of PCBP1-mediating critical interactions on viral replication and helping to develop better antiviral solution.

## Introduction

Classical swine fever (CSF), driven by CSF virus (CSFV), is a highly contagious porcine disease, causing substantial economic losses^[1, 2]^ and the typical clinical signs are generally characterized by high fever, inappetence, and general weakness followed by neurological deterioration, skin hemorrhages, and splenic infarction^[3, 4]^. The genome of CSFV could encode four structure proteins (C, E^rns^, E1, and E2) and eight non-structure proteins (N^pro^, p7, NS3, NS4A, NS4B, NS5A, and NS5B), which would utilize host factors for enhancing replication and evading cellular immunity^[5]^. It has been confirmed that envelope protein E^rns^ would interact with HS or LamR for the attachment of CSFV particles to the surface of permissive cells and that structure protein E2 interacted with Anx2 and/or MEK2 to promote CSFV production^[6]^. Recently, it was proposed that N^pro^ could interact with a host factor designated as PCBP1 which is positive for CSFV replication^[7]^.

Poly (rC)-binding protein 1 (PCBP1), an RNA- or DNA-binding protein, could regulate the process of pre-mRNA, mRNA stability, and translation in nature^[8, 9]^. It also participated in the formation of iron chaperone complex, influencing the delivery of iron in cell^[10]^. Additionally, deficiency of PCBP1 could decrease the apoptosis induced by heavily oxidized RNA in human cells^[11, 12]^. On the other hand, in the virus-host interplay area, it was suggested that PCBP1 was associated with cGAS in a viral infection-dependent manner and promoted cGAS binding to DNA. PCBP1 deficiency inhibited cytosolic DNA- and DNA virus-triggered induction of downstream effector genes^[13]^. Moreover, PCBP1 could mediate housekeeping degradation of MAVS via ubiquitination by a E3 ubiquitin ligase called AIP4 and overexpression of PCBP1 inhibited SeV-induced antiviral responses^[14]^. Although the PCBP1 is conserved across various species, due to the reason that retrotransposition of it from a processed PCBP2 predates the mammalian radiation^[15, 16]^, the function of it may be divergent, especially in the duration of virus infection. It has been reported that PCBP1 interacted with PRRSV nsp1β and colocalized with viral replication and transcription complex (RTC) ^[17]^, but the confirmation was performed in Marc-145 cell line which was not porcine cells. What the definite roles of PCBP1 in cells or individuals of porcine origin in the duration of viral infection is needs to be further investigated.

Although vaccines have been widely used to control CSFV infections in population, sporadic individuals occurred continuously^[5, 6, 18]^. To fundamentally counteract with the consequence caused by CSFV, more effective and endogenous strategies are needed to be adopted. Genetic modification in pigs is one of efficacious strategies that has been adopted to generate pigs with resistance to various swine viruses, such as PRRSV^[19, 20]^, TGEV^[21]^ using CRISPR/Cas9 technology. Hence, based on the host factors hijacked by corresponding viruses which play critical roles in viral entry, internalization, and replication, creating pigs with viral resistance via knockout method is promising.

Herein, we knock out *PCBP1* gene in PK-15 cells as well as primary porcine fibroblasts (PFFs) using CRISPR/Cas9 technology and characterize the anti-CSFV ability of *PCBP1* KO cell clones. Meanwhile, we generate *PCBP1*^*-/+*^ pigs through somatic cell nuclear transfer (SCNT) with *PCBP1* KO PFFs. Additionally, the effect of PCBP1 deficiency on the IFN-α pathway and predicted interactors of PCBP1 after CSFV infection was also explored.

## Results

### Generation of *PCBP1* knockout PK-15 cells

First of all, the *PCBP1* relative expression in various porcine organs was detected (Fig. 2a). Within N terminus of the only exon in PCBP1 locus, two 20-base-pair (bp) sequence were selected (Fig. 2b). Based on both crRNA sequence, pX330 plasmids expressing different guide RNAs were created which were designated as sg97 and sg95 respectively (Fig. 2b). The cleavage efficiency of both sgRNAs were monitored via transient electrotransfection into PK-15 cells (Fig. 1a). As shown in Fig. 2c and 2d, although the efficiency of sg97 was slightly higher than that of sg95, both of them were allowed to participate in following investigation.

**Fig. 1.**
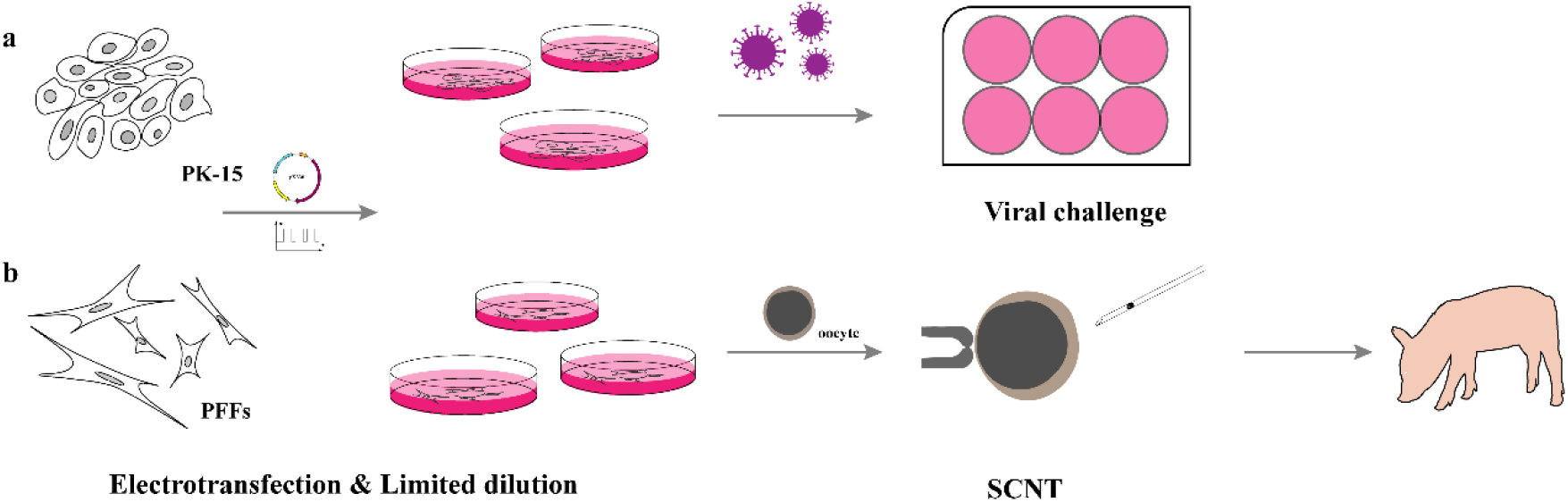
The overall design of this study. (**a**) The screen of sgRNA with high efficiency and the selection of PK-15 positive clone, as well as viral challenge assay in vitro. (**b**) The circuit of generation of gene-editing piglets.

**Fig. 2.**
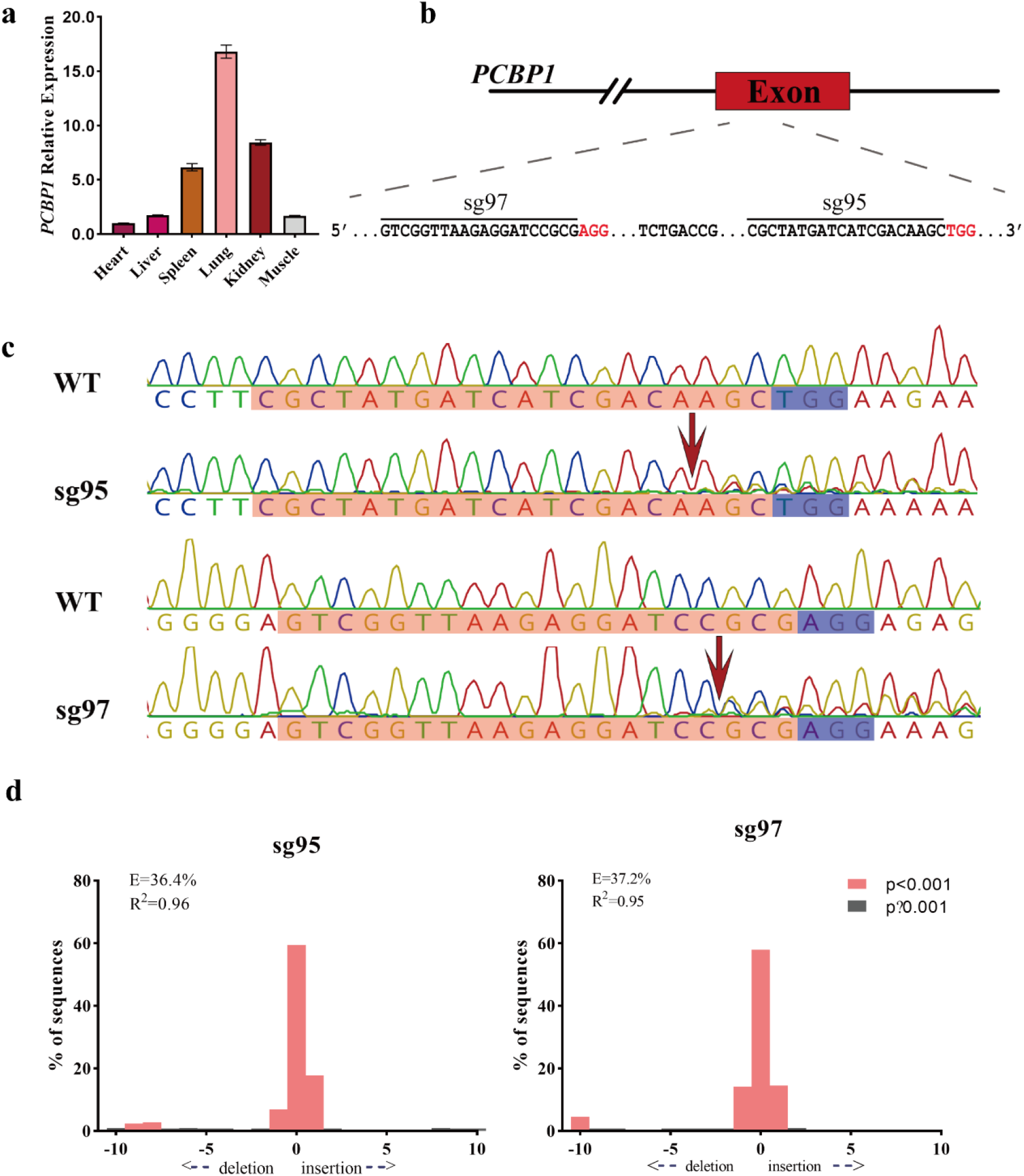
The screen of sgRNA. (**a**) The relative expression level of *PCBP1* in various organs from Large White piglet was determined by RT-qPCR. (**b**) The targeting diagram of representative sgRNAs on *PCBP1* locus. The red bases indicate PAM sequence. (**c**) The corresponding cutting efficiency of sgRNAs in *b* is analyzed by Sanger sequencing. The red arrow indicates the cleavage site of Cas9 protein. The bases in purple rectangle are PAM sequence. The bases in orange rectangle are crRNA sequence. (**d**) The cleavage efficiency of corresponding sgRNAs in *b* are visualized using TIDE.

To select and identify *PCBP1* KO clones, sg97 and sg95 were separately electrotransfected into PK-15 cells. *PCBP1* KO positive clones were selected with limited dilution method. Total 49 clones were detected and 5 positive clones were obtained. As shown in Fig. 3a, a subset clones were examined through T7 endonuclease I (T7E1) assay in which 15#, 25#, and 27# were sg97-producing positive clones and 40# and 46# were sg95-producing positive clones. To verify the genotype of positive clones, we performed T-cloning and Sanger sequencing using specific primers amplifying segment containing sgRNA-targeting region. Three positive *PCBP1* KO clones were chosen to perform further research. Ten bp proximal to PAM sequence were deleted in 15# clone and 1 bp was deleted in 27# which were compound heterozygous *PCBP1* KO clones. Otherwise, as for sg95-producing positive 40#, homozygous clone, a T and a A were respectively inserted into each chromosome locus (Fig. 3b).

**Fig. 3.**
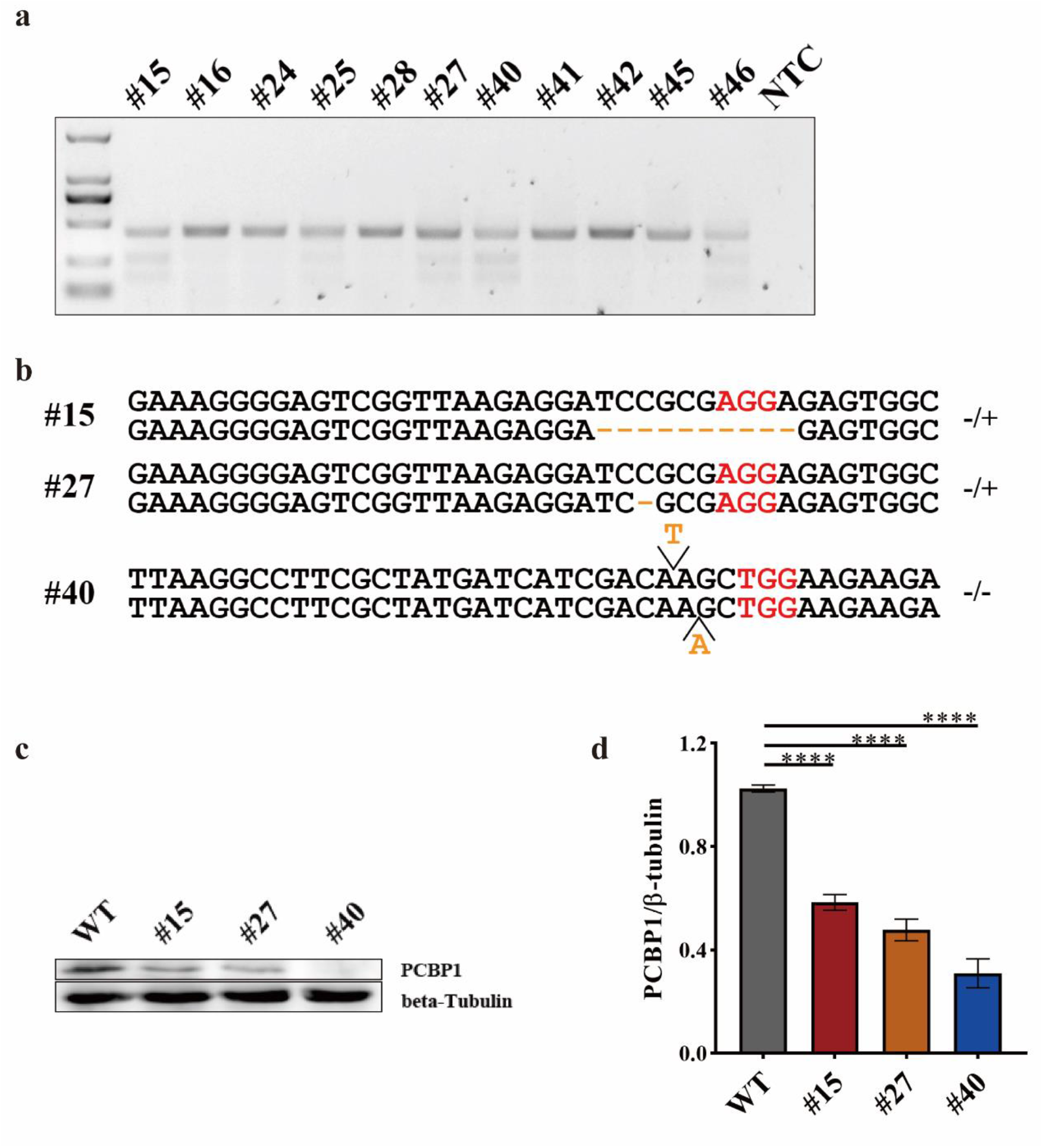
The screen of *PCBP1* KO clones in PK-15 cells. (**a**) The cleavage efficiency in various selected clones are detected by T7E1 cleavage assay. M, DL2000, has been used to indicate band size. (**b**) T-cloning and Sanger sequencing of editing *PCBP1* alleles in different type of positive clones. PAM sites are highlighted in red. Indels are shown in yellow. (**c**) Endogenous PCBP1 level of various positive KO clones was determined by western blotting. (**d**) The gray intensity analysis of PCBP1. PCBP1 band intensity was normalized to that of beta-tubulin in the same sample. Every sample was measured three times by ImageJ. Bars are presented as mean ± SEM and data are analyzed by Student’ s *t*-test using Graphpad Prism 8.0. *p < 0.05; **p < 0.01; ***p < 0.001; ****p < 0.0001; n = 3.

To confirm the loss of PCBP1 expression in above selected positive clones, western blot was performed. As shown in Fig. 3c, PCBP1 deficiency occurred not only in homozygous KO clone (40#) but also in heterozygous clones (15# and 27#) in comparison with the wild type PK-15. Eventually, gray intensity value analysis of corresponding band also indicated that PCBP1 level in KO clones was notably reduced compared to that in WT cells. These data above demonstrated that *PCBP1* was successfully knocked out in PK-15 cells and several positive KO clones were obtained.

### *PCBP1* KO PK-15 cells inhibit CSFV proliferation but not PRV and PEDV

To explore the antiviral capability of *PCBP1* KO positive clones, *PCBP1* KO clone Number 15 and clone Number 40 were infected by several swine viruses. It is reported that knockdown of PCBP1 could suppress CSFV growth^[7]^. Hence, quantitative reverse transcription PCR (RT-qPCR) was performed to detect the number of viral genomes at various hours post-infection (hpi) firstly. The magnitude of CSFV genome was significantly reduced in #15 and #40 compared to WT from 12 hpi to 48 hpi (Fig. 4a). This finding coincided with the immunofluorescence assays showing that the expression of the CSFV-encoded E2 protein in PCBP1 KO cells was reduced following CSFV infection (Fig. 4d). The fluorescence intensity indicated that viral load in PCBP1 KO clone was less than that in WT (Fig. 4e). Meanwhile, the magnitude of CSFV genome in corresponding time point was consistent with IFA result (Fig. 4f). Then, PEDV and PRV challenge were also performed comparable with CSFV. However, the level of viral genomes in PCBP1 KO clones was consistent with WT for PEDV (Fig. 4b) and PRV (Fig. 4c). The viral load of PRV at various time point was chaotic probably due to the cytopathic effect (CPE). Taken together, these results suggest that PCBP1 knock out could significantly inhibit CSFV growth in PK-15 cells.

**Fig. 4.**
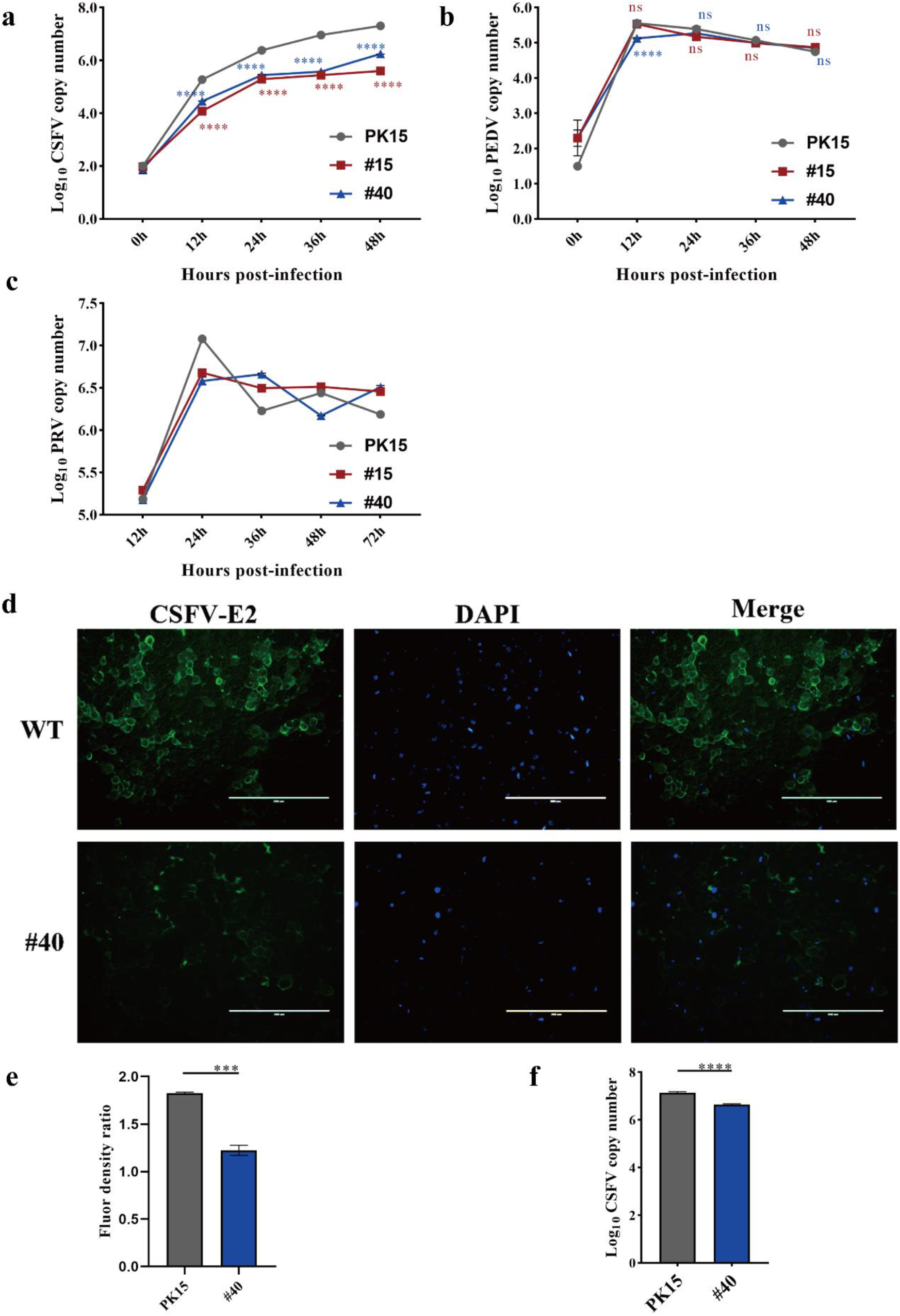
PCBP1 knockout could reduce CSFV infection but not PRV and PEDV. (**a**)The proliferation kinetics of CSFV in PCBP1 KO positive clones at various time points post-infection. (**b**) The proliferation kinetics of PEDV in PCBP1 KO positive clones at various time points post-infection. (**c**) The proliferation kinetics of PRV in PCBP1 KO positive clones at various time points post-infection. (**d**) Viral resistance to CSFV was examined by IFA. (**e**) The mean fluorescence intensity in d was analyzed by ImageJ. (**f**) The copy number of CSFV genome at the same hpi with d was detected by RT-qPCR. Bars are presented as mean ± SEM and data are analyzed by Student’ s *t*-test using Graphpad Prism 8.0. *p < 0.05; **p < 0.01; ***p < 0.001; ****p < 0.0001; ns, no significance; n = 3.

### PCBP1 knockout potentiates innate antiviral responses stimulated by CSFV in PK-15 cells

To further investigate the mechanism of inhibition for CSFV but not for PEDV in PCBP1 KO cell line, we detected the relative expression level of several type I interferon (IFN) genes, such as IFN-alpha and IL-6 prior to the ISGs in PCBP1 KO cells. Clone 40# was chosen as following objective cell line. Compared to that in PEDV-infecting groups, the transcription level of IFN-alpha and IL-6 in CSFV-infecting groups was increased around 8-fold and 2-fold respectively (Fig. 5a). Progressively, the transcription level change of various interferon-stimulated genes that have antiviral activity against a board range of viruses was further explored. As shown in Fig. 5b, the relative expression of effector genes, downstream genes of interferon, were universally higher than that in PEDV-infecting cells.

**Fig. 5.**
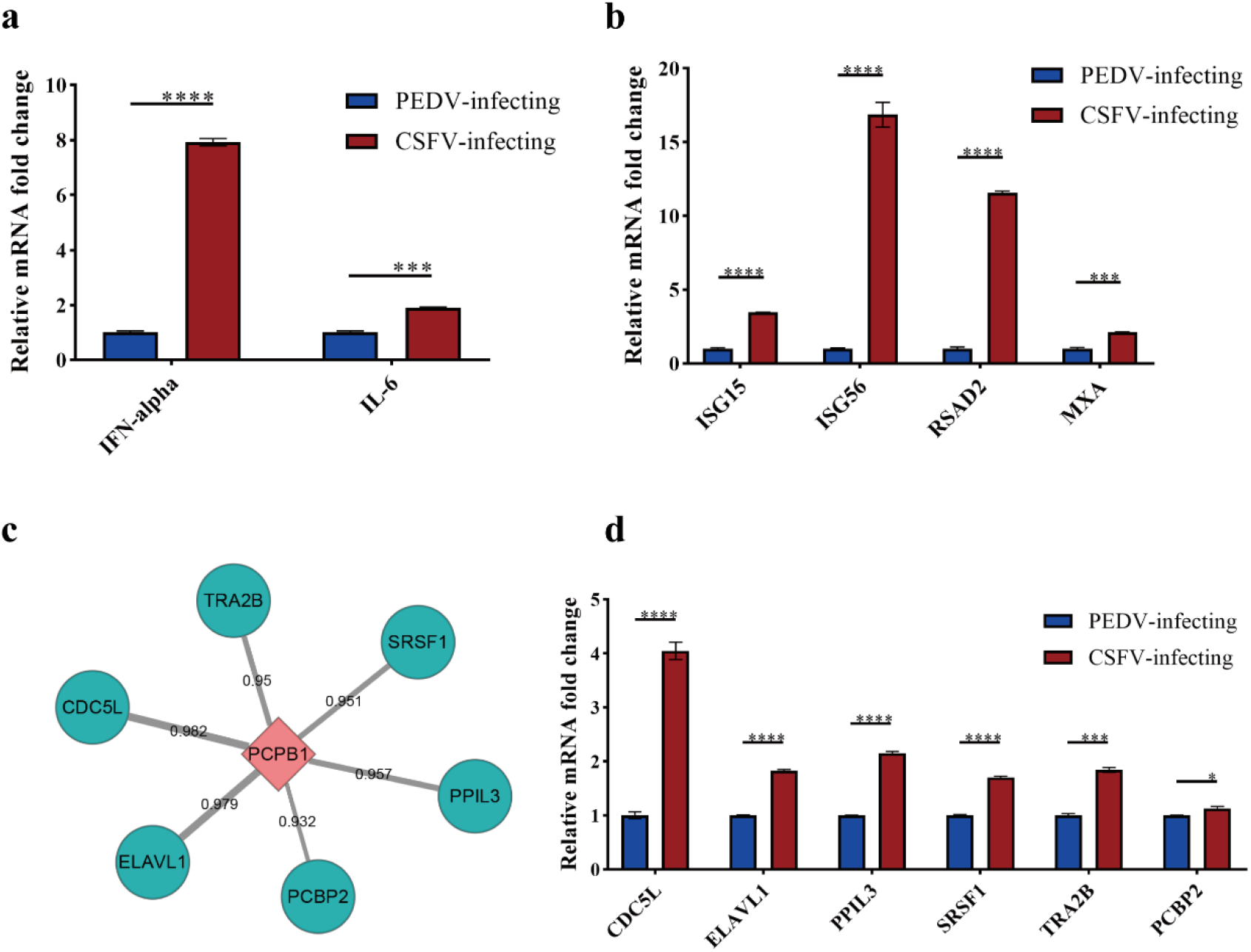
The alteration of IFN associated effectors and predicted genes related to PCBP1. The relative mRNA fold change of IFN pathway genes (**a**) or the downstream effectors (**b**) assessed in *PCBP1*^*-/-*^ PK-15 clone using RT-qPCR at 36 h postinfection. (**c**) The predicted interactors of PCBP1. The thickness of the gray line represents combined score. (**d**) The relative mRNA fold change of predicted genes assessed in *PCBP1*^*-/-*^ PK-15 clone using RT-qPCR at 36 h postinfection. PEDV-infecting samples were used as reference samples. Bars are presented as mean ± SEM and data are analyzed by Student’ s *t*-test using Graphpad Prism 8.0. *p < 0.05; **p < 0.01; ***p < 0.001; ****p < 0.0001; ns, no significance; n = 3.

Additionally, in order to observe the alteration of interplay relative to PCBP1, we searched for the interactors of PCBP1 using STRING database^[22, 23]^, and the top six predicted genes were shown in Fig. 5c. Interestingly, all of these predicted genes were more up-regulated in CSFV-infecting 40# clone than PEDV-infecting samples. Taken together, the cytokines of innate immunity induced by CSFV in PCBP1 KO cells were more intensive than that stimulated by PEDV.

### Primary fibroblasts derived from PCBP1 KO pigs diminish CSFV infection

Our major goal in this research was to generate a herd of *PCBP1* KO pigs, which could inhibit CSFV infection. To achieve this purpose, *PCBP1* KO PFFs should be produced firstly (Fig. 1b). The sg97 were introduced into Large White PFFs and the positive clones were selected comparable with the operation in PK-15 cells. Prior to SCNT, cell viability of *PCBP1* KO PFFs were monitored by CCK8. As shown in Fig. 6a, knockout of *PCBP1* in PFFs did not exert notable adverse effects. The *PCBP1* KO PFF clone was used as donor cells for SCNT and total 921 matured reconstructed embryos were transferred into five surrogates. The piglets were born after around 114 days of pregnancy, two of which were shown in Fig. 6b and three of them were identified as positive heterozygous *PCBP1* KO pigs by PCR and Sanger sequencing (Fig. 6c). To elucidate the effect of knockout on genome of F_0_ pigs, off-target sites located on different chromosomes were predicted using RGEN tools and no obvious off-target events occurred as shown in Fig. 7a and 7b.

**Fig. 6.**
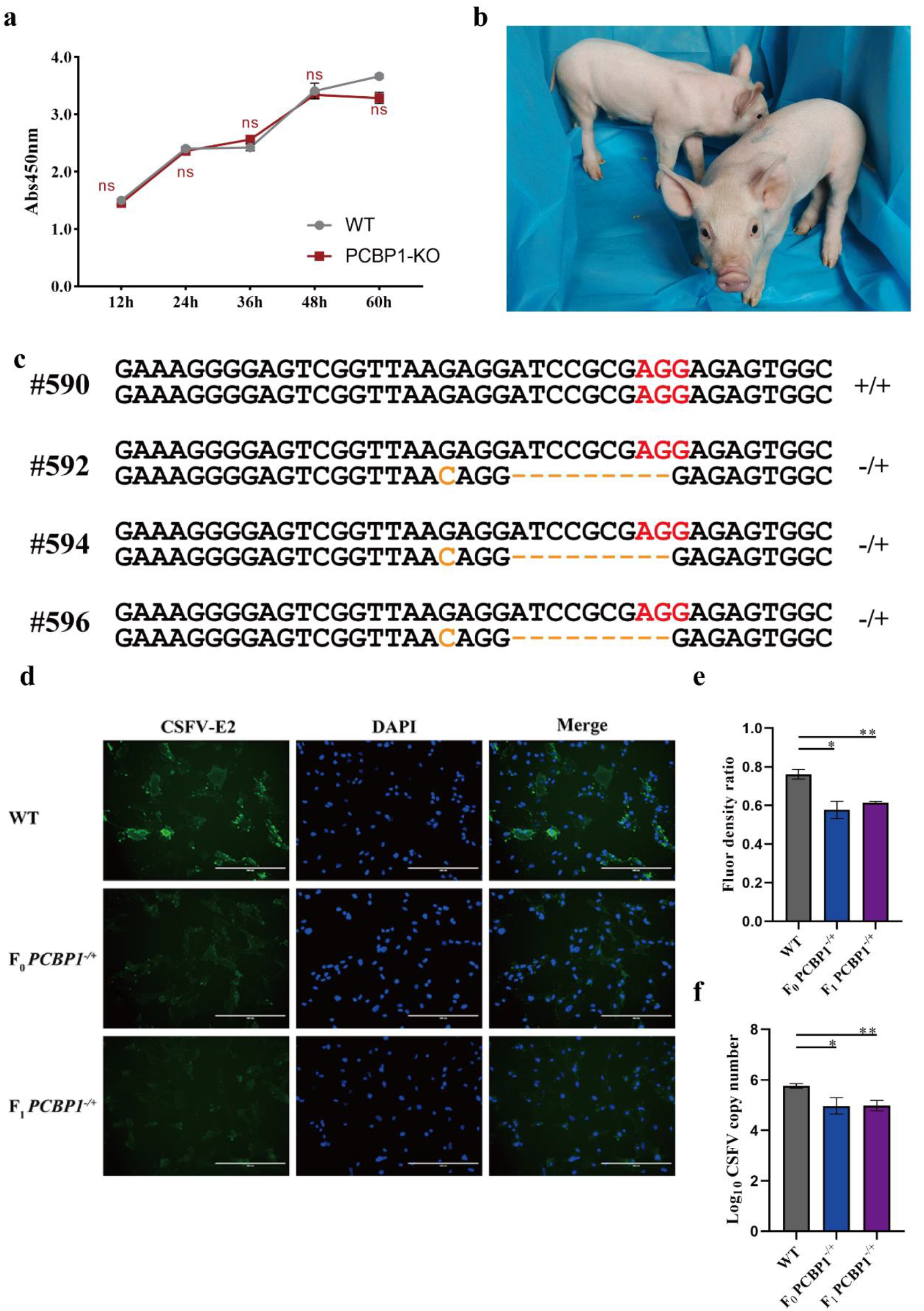
Production of *PCBP1*^*-/+*^ pig. (**a**) The cell viability of PCBP1 KO PFFs. (**b**) Photograph of F_0_ *PCBP1*^*-/+*^ piglets. (**c**) T-cloning and Sanger sequencing of *PCBP1* alleles in F_0_ piglets. (**d**) The anti-CSFV ability of F_0_ and F_1_ pigs was detected using primary tail fibroblasts by IFA. (**e**) The mean fluorescence intensity in *d* was analyzed by ImageJ. (**f**) Genomic replication of CSFV in primary tail fibroblasts of F_0_ and F_1_ pigs was detected by RT-qPCR at 36 hpi. Bars are presented as mean ± SEM and data are analyzed by Student’ s *t*-test using Graphpad Prism 8.0. *p < 0.05; **p < 0.01; ***p < 0.001; ****p < 0.0001; ns, no significance; n = 3.

**Fig. 7.**
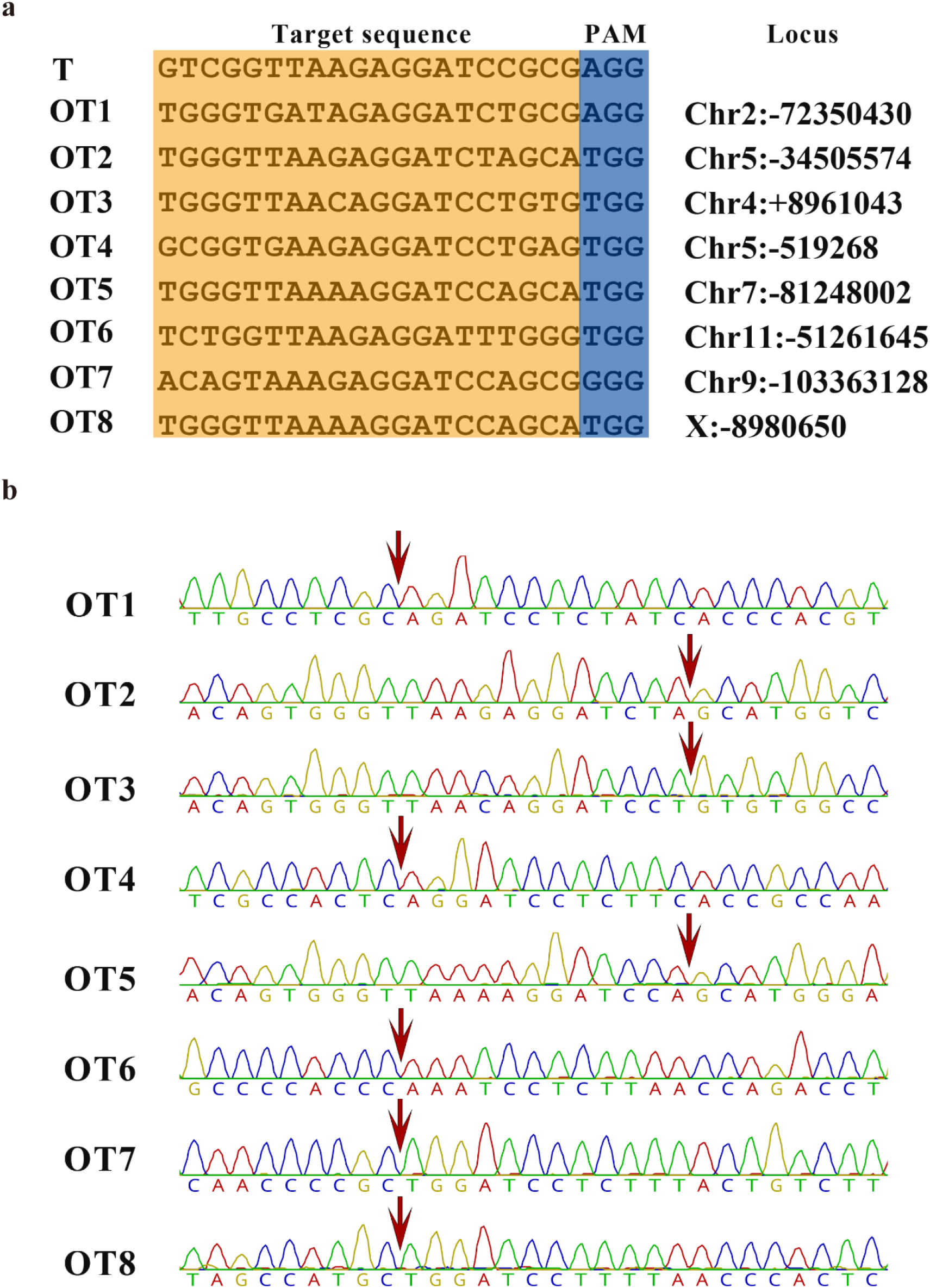
Off-target analysis. (**a**) The target site (T) and eight predicted off-target sites (OT) of sg97. OT1∼OT8 indicates eight off-target sites and T represents target site. (**b**) Sanger sequencing results of PCR amplicons of each off-target site. The red arrow indicates the potential cleavage sites.

To expand the herd of *PCBP1* KO pigs, the female of F_0_ generation mated with wild type herd boar while the positive pigs grown to the estrus period. Recently, the offspring of F_0_ generation was born and the alleles of *PCBP1* were also confirmed by Sanger sequencing as above. In order to verify the anti-CSFV ability, primary fibroblasts isolated from tail tips of *PCBP1*^-/+^ F_0_ and F_1_ were infected by CSFV for 36 h. As shown in Fig. 6f, the magnitude of CSFV in *PCBP1* KO PFFs was significantly decreased in comparison with that in WT. The similar result was further indicated by IFA (Fig. 6d and 6e). Altogether, we prepared *PCBP1*^-/+^ pigs and expanded the herd of it, which had the potential to inhibit the proliferation of CSFV.

## Discussion

CSFV, the pathogen of CSF which is characterized by multiple hemorrhages, leukopenia, high fever, abortion, and neurological dysfunction^[1, 24]^, has caused significant economic losses worldwide. Production of transgenic pigs is one of the powerfully effective strategies to contain viral infection by alteration of immune state genetically which has been widely utilized to resist various porcine viruses^[25-28]^. While different types of genetically modified pigs were generated via exploiting the key host factors responding to viral infection^[29, 30]^, the pigs of endogenous restricted factors knockout rarely occurred. In this reported, we targeted the PCBP1 locus in porcine genome using CRISPR/Cas9 technology and successfully acquired *PCBP1* KO PK-15 cell line and *PCBP1*^*-/+*^ individual pigs. In vitro and ex vivo viral challenge both illustrated that *PCBP1* KO cells could significantly reduce CSFV infection. To the best of our knowledge, this study is the first report of *PCBP1* knockout pigs with the resistance to CSFV.

It was proposed that heterozygous *PCBP1* in mouse displayed a mild and nondisruptive defect in initial postpartum weigh^[15]^. However, the F_0_ generation of *PCBP1*^*-/+*^ pigs exhibited normal birth weight and phenotype which may demonstrate that PCBP1 plays divergent roles in the duration of development between mouse and pigs. A previous report indicated that overexpression of PCBP1 could enhance CSFV growth and reasoned that the deletion of KHIII would cause PCBP1 incorrect folding, leading to abrogation of PCBP1-N^pro^ interaction^[7]^. Profressively, we provide a possible hypothesis that the precise amino acid residue position or positions which play an important role in the interaction with N^pro^ may locate on the KHIIII domain. The base editing library screen derived from CRISPR/Cas9 technology is developing with a high speed and has been widely used^[31-33]^. Comprehensive screen of the precise amino acid in PCBP1 with saturation editing is perspective for addressing specific sites interacting with N^pro^ and exploitation of targeted drugs.

Type I interferon (IFN) has antiviral activity and RNA viruses of the family Flaviviridae are sensitive to type I IFN^[5, 34]^. Besides, Activation of type I IFN could induce synthesis of hundreds of proteins such as interferon-stimulated genes (ISGs) ^[34, 35]^. It was suggested that CSFV N^pro^ was involved in inhibition of type I IFN by interaction with IRF3^[36, 37]^. Our data demonstrated that type I IFN genes and the downstream ISGs such as ISG15, ISG56, and RSAD2, all of which were well-documented to inhibit a broad spectrum of viruses^[35, 38-41]^, were increased following CSFV infection in PCBP1 deficient PK-15 cells, implying the enhancement of cellular innate immunity. In terms of the literature above and our results, we speculate that PCBP1 may involve in the process of N^pro^ inhibition against type I IFN. In undisturbed infection states, PCBP1 participates in the conformation of N^pro^-IRF3 complex to suppress type I IFN induction and the activation of downstream effectors. However, deficiency of PCBP1 blocks the conformation of N^pro^-IRF3 complex, which limits the reduction of type I IFN cascade reaction. A previous report illustrated that knockdown of PCBP1 promoted the increase of type I IFN in cells infected with SeV or transfected with poly (I:C)^[14]^, which confirmed our speculation to a certain degree. However, the function of PCBP1 in process of CSFV counteracting cellular immune system infection is still unclear. Differently, it is reported that type III IFNs play critical roles in innate antiviral immunity in intestinal epithelial cells in the gut^[42, 43]^. We reason that the depletion of PCBP1 do not influence the immune responses following PEDV infection because PCBP1 may not be included in type III IFN cascade reaction.

To further explore the post alteration of relevant genes due to deficiency of PCBP1 in the presence of CSFV, we predicted the interactors of PCBP1 using STRING database. Among the predicted genes, it was proposed that overexpression of several interactors such as ELAVL1 and SRSF1 would decrease the level of adenovirus, ZIKV, and HIV-1, etc.^[44-47]^, implying the property of these genes of inhibiting viral infection. Our results illustrated that CDC5L, ELAVL1, and SRSF1, etc. were universally upregulated after CSFV stimulation which may be restricted factors relative to PCBP1 in the duration of CSFV infection. PCBP1 hijacked by N^pro^ or other CSFV proteins suppress the activation of some antiviral pathways including above detected factors. The removal of inhibition leads to the upregulation of ELAVL1 and SRSF1, etc. following CSFV infection due to the deficiency of PCBP1.

Recently, the PCBP1 knockout pigs of F_1_ generation, offspring of heterozygous 592#, was successfully produced. As expected, ex vivo cultured primary cells isolated from F_1_ generation still displayed significant anti-CSFV capability. Unfortunately, the first litter was so small that the following research cannot be performed. The herd of *PCBP1* KO pigs is strictly monitored until the scale of research recipients is large enough to execute following experiments and individual level schedule concerning in vivo viral challenge is preparing now. In further future, the ex vivo results will be directly translated into in vivo model promisingly.

In summary, the PCBP1 knockout pigs are not only a valuable animal model for further investigating infection mechanisms of CSFV but also hold the potential to reduce economic losses related to CSFV in swine industry.

## Materials and Methods

### Cell Lines and Culture Conditions

Porcine kidney cell line-15 (PK-15) cells (ATCC Number: CCL-33) were cultured in Dulbecco’s modified Eagle’s medium (DMEM, Gibco) supplemented with 5% fetal bovine serum (FBS), 10 Unit/mL penicillin, 10 μg/mL streptomycin, 1% Non-Essential Amino Acids (NEAA, Gibco), and 2 mM L-Glutamine (Gibco). Primary porcine fatal fibroblasts (PFFs) were cultured in DMEM containing 15% FBS, 10 Unit/mL penicillin, 10 μg/mL streptomycin, 1% NEAA, and 2mM L-Glutamine. All cells were grown in an atmosphere of 5% CO_2_ at 37°C.

### Viruses

CSFV Shimen strain and PRV (Suid herpesvirus 1) were used and maintained at -80 °C. PEDV attenuated vaccine was purchased from Jilin Zhengye Biological Products CO., LTD. All attenuated virus in dry powder form was stored at 4°C and the stock solution was preserved at -80°C.

### Plasmid Construction

CrRNA sequence was searched through the porcine *PCBP1* gene using the CHOPCHOP webtools (https://chopchop.cbu.uib.no/). CACC sequence was added at 5’ end of the top strand of selected crRNA sequences and AAAC was added at 5’ end of the bottom strand. These sgRNA oligonucleotides were synthesized by Comate Bioscience CO., LTD and ligated into the Bbs I sites of pX330 vector (42230, Addgene) to form the intact targeting plasmids.

### Electroporation and Generation of Knock Out Cell Clones

Approximately 30 μg pX330 plasmids containing crRNAs targeting different region of porcine PCBP1 gene were electrotransfected into ∼3 × 10^6^ PFFs using Neon Transfection System (invitrogen). The specified parameters applied to PFFs uniquely were as follows: 1260 voltage, 30ms, 1 pulse. Similarly, 30μg pX330 plasmids were introduced into ∼3 × 10^6^ PK-15 cells resuspended in 300μL Opti-MEM (Gibco) in 2 mm gap cuvettes using BTX-ECM 2001. The parameters were as follows: 300 voltage, 1 ms, 3 pulses, 1 repeat. The PFFs and PK-15 cells were seeded into ten 100mm dishes after 48 hours post-transfection, and the inoculation density per dish was 2000 cells on average. The cell clones were picked and continually cultured in 24-well plates. Forty percent cells per well were digested for 2 min at 37°C and lysed with 10μL NP-40 lysis buffer (10 mM Tris-HCl pH 8.3, 50 mM KCl, 1.5 mM MgCl_2_, 1% NP-40, and 1% protease K) for 1 h at 56°C and 10 min at 95°C after each clone reaching into 80% confluency. The lysate was used as PCR template and subjected to Sanger sequencing. The positive PK-15 clones were propagated into 100 mm dishes one step at a time. The positive PFFs clones were grown on 24-well plates until SCNT.

### T7E1 assay

Genomic DNA of positive PFFs clones was extracted using TIANamp Genomic DNA Kit (TIANGEN). And a conventional PCR was performed as follows: 95°C for 4 min; 95°C for 30 s, 59°C for 30s, 72°C for 30s, for 35 cycles; 72°C for 5 min; hold at 4°C. The PCR products were purified using QIAquick PCR Purification Kit (Qiagen). Approximately 200 ng purified PCR products mixed with 10 × NEB

Buffer 2 were hybridized using following cycles: 95°C for 5 min; 95-85°C at the rate of -2°C/s, 85-25 at the rate of -0.1°C/s; hold at 4°C.Then, 1 μL T7 endonuclease was added to each sample and the reactions were incubated at 37°C for 15 min. the reaction mixtures were then analyzed on a 2% agarose gels.

### Virus infection

The in vitro viral challenge assay was stringently performed and monitored at a designated safe place. The positive clones or primary PPFs were seeded in 6-well plates. For CSFV and PRV infection, cells were replaced with fresh culture medium after incubating for 1 h at a multiplicity of infection (MOI) of 20 and 50 respectively. For attenuated PEDV infection, the absorption phase was maintained for 1 h at a MOI of 10 in the presence of 10 μg/mL trypsin, after which the maintenance medium containing 10 μg/mL trypsin was added. At various time points postinfection, samples containing viral genome were harvested and stored at -80°C until use.

### Viral genome extraction and Real-Time quantitative PCR

As for CSFV, total cellular RNA was extracted from CSFV-infecting PK-15 cells or positive clones using TRNzol Universal Reagent (TIANGEN) and ∼2 μg RNAs were performed to reverse transcript to the first-strand cDNAs using FastKing RT Kit (TINAGEN) according to manufacturer’s instruction. As for PEDV and TGEV, the monolayer of virus-infected cells were scraped by cell scraper within the culture medium and 200 μL suspension was aspirated and mixed with 800 μL TRNzol Universal Reagent. The subsequent reverse transcription was consistent with the above. As for PRV, the virus-infected material was obtained in the same manner as PEDV and TGEV. And the PRV genome within 200 μL suspension was extracted by TIANamp Virus DNA/RNA Kit (TIANGEN). All cDNAs and viral genome were -20°C.

To detect the accurate viral copy number in virus-infected materials, a standard curve was generated with 10-fold serial dilutions ranging from 10^9^ to 10^3^. The quantitative PCR was performed using Quantagene q225 (KUBOTECHNOLOGY) with SuperReal PreMix Plus (TIANGEN) according to the manufacturer’s instruction. To check the relative expression of predicted genes or genes associated with porcine PCBP1, the housekeeping gene glyceraldehyde 3-phosphate dehydrogenase (*GAPDH*) was selected as reference gene and the mRNA expression was normalized to *GAPDH* using the 2^-ΔΔCt^ method.

### Western Blotting

The wild type PK-15 and cell clones were washed in ice-cold phosphate-buffered saline (PBS) and lysed in Cell Lysis Buffer for Western And IP (P0013, BEYOTIME) in the presence of 1mM PMSF (AR1192, BOSTER) and 1% Protease Inhibitor Cocktail (P1005, BEYOTIME). The protein concentrations were measured with the BCA assay Kit (AR1189, BOSTER) and 40 μg proteins were diluted in 5 × SDS-PAGE Loading Buffer (AR1112, BOSTER) at 95 °C for 10 min. Subsequently, the samples boiled were resolved on the artificial 4∼12% SDS-PAGE gel and the proteins were transferred to nitrocellulose membranes. The membranes were blocked with 5% skim milk dissolved in TBST for 2 h at room temperature. Primary antibodies for immunoblotting were as follows: rabbit anti-PCBP1 (1:2000, BOSTER A02636-1), rabbit anti-β-tubulin (1:5000, BOSTER BM3877). Membranes were subsequently washed in TBST and then incubated with horseradish peroxidase-conjugated goat anti-rabbit/mouse IgG (H + L) (1:5000, BOSTER BA1056). Ultimately, membranes were imaged with the ultra-sensitive ECL chemical luminescence ready-to-use kit (BOSTER AR1197) using Azure c600 (AZUREBIOSYSTEMS). The corresponding protein bands were normalized to β-tubulin band density using Fiji.

### IFA

The positive clones or primary fibroblast cells isolated from tail tips of the *PCBP1*^*-/+*^ F_0_ piglets were seeded into 24-well plates with four replicates per sample. The cells, reaching 80% confluency, were infected with CSFV (200 TCID_50_ per well). At 2 h post-inoculation, cells were replaced with fresh CSFV-free culture medium. After 36h inoculation, cells were washed with cold PBS and fixed in 4% paraformaldehyde for 30 min at room temperature. The primary antibodies and fluorophore-conjugated antibody were as follows: mouse anti-CSFV E2 (1:500, LVDU BIO-SCIENCES & TECHNOLOGY CO., LTD.), fluorescein (FITC)-conjugated goat anti-mouse IgG (H + L) (1:500, PROTEINTECH SA00003), and Alexa Fluor 488-conjugated goat anti-mouse IgG (H + L) (1:500, PROTEINTECH SA00006). Samples were incubated with primary antibodies for 1 h in cold blocking buffer (10%FBS in PBS) at 37°C, followed by three washes in PBS and incubated with secondary antibodies in a dark, humidified chamber for 1 h at 37°C. Before imaged with EVOS f1 fluorescence microscope, samples were washed five times with PBS. The semi-quantitative fluorescence intensity of the target protein was normalized to that of corresponding nucleus using Fiji.

### SCNT

The *PCBP1*^*-/+*^ positive PFFs were used for somatic cell nuclear transfer as described previously^[48]^. The positive cells were injected into the perivitelline cytoplasm of enucleated oocytes to form reconstructed embryos. Subsequently, reconstructed embryos were surgically transferred into the oviducts of surrogate females on the first day of estrus after activated and cultured for approximately 18 h in embryo culture medium. Pregnancy status was detected using ultrasound scanner between 30–35 days post-transplantation. To monitor the blastocyst formation rate and developmental viability, a part of activated embryos was continually cultured for 7 days. **Isolation of primary porcine fibroblast**. The tail tips from *PCBP1*^*-/+*^ and WT piglets were cut into small pieces, followed by digested with the fresh culture medium containing 20% FBS in the presence of 25 Unit/mL DNase I and 200 Unit/mL type IV collagenase for 4 h at 39 °C. Then, dissociated primary cells and tail pieces were continually cultured for 4∼5 days. The isolated tail fibroblasts were cryopreserved at - 80 °C for 24h, after which moved to liquid nitrogen for long term storage.

### Cell viability assay

cell viability was evaluated with the Cell Counting Kit-8 (AR1160, BOSTER) according to the manufacturer’s instruction. Briefly, the PCBP1 KO PFFs or WT cells were seeded into 96-well plates at a density of 5 × 10^3^ cells/well. The cells were replaced with fresh culture medium containing 10% CCK-8 reagent until attached to plates. An additional inoculation were applied for 1 h at 37°C. The absorbance at 450nm was measured using TECAN Infinite 200 PRO.

### Statistical analysis

Statistical analysis was performed using Graphpad Prism 8.0 software. Student t tests were used to compare two groups. *P* < 0.05 was considered statistically significant.

## Acknowledgments

This work was financially supported through grants from the Special Funds for Cultivation and Breeding of New Transgenic Organisms (No. 2016ZX08006003) and the Shenzhen Key Technology Projects (JSGG20180507182028625).

---

**Table 1.**
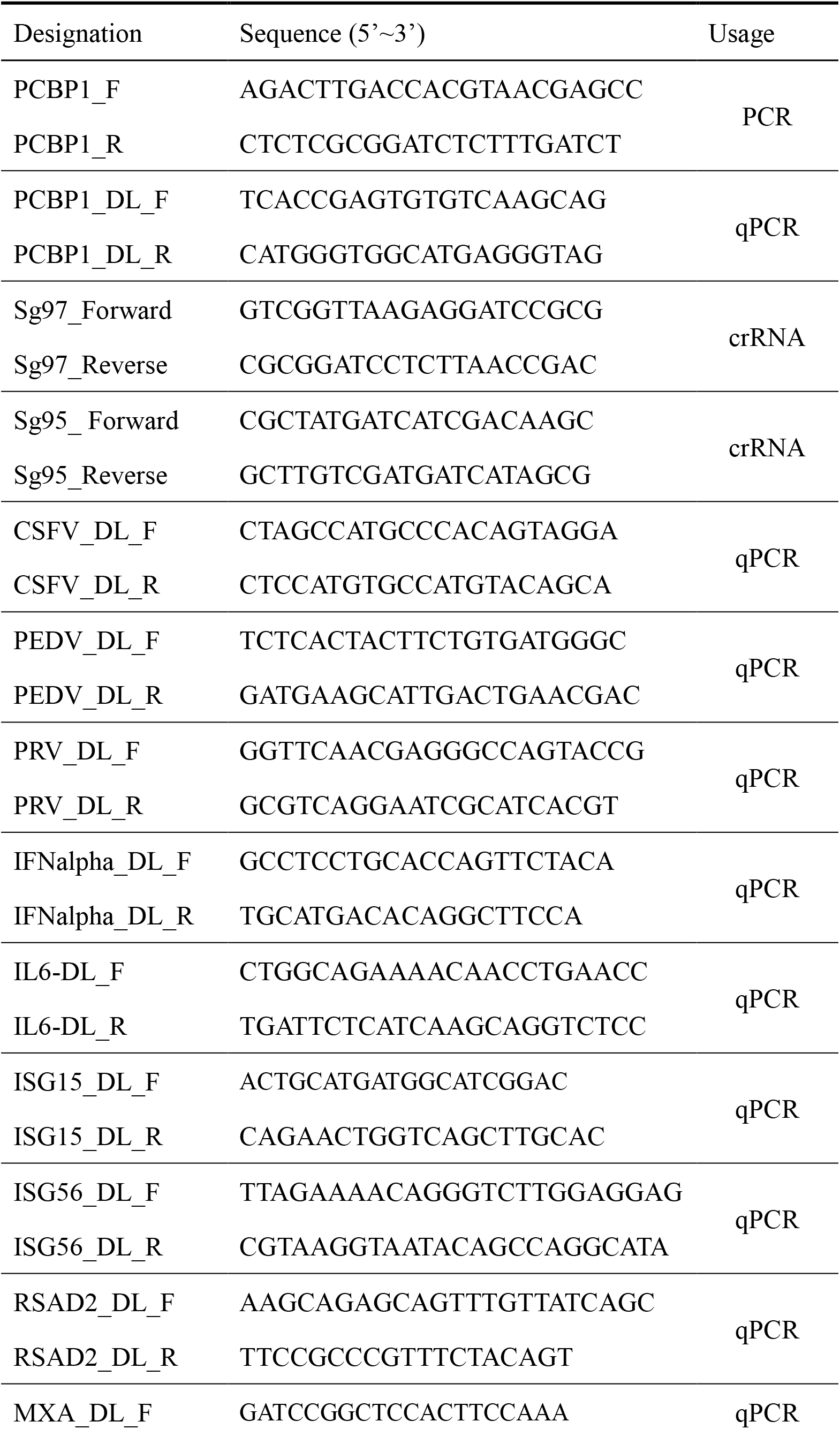
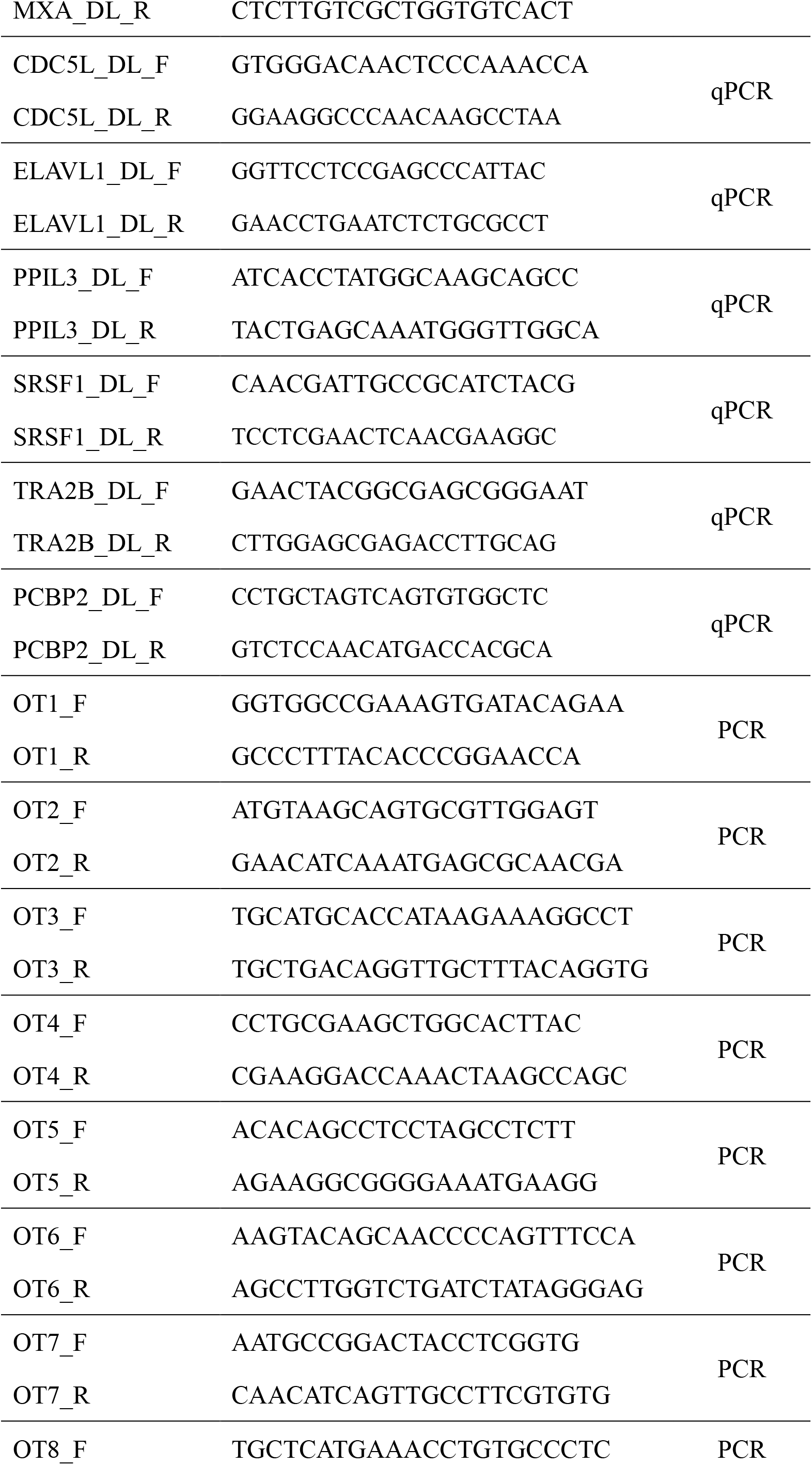
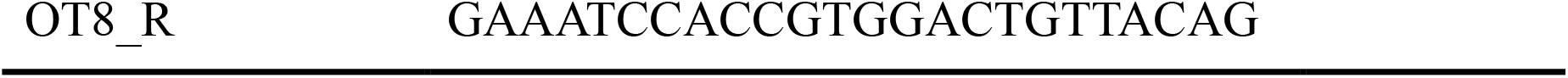
Primers and sequences in this research

